# Fully automated, sequential focused ion beam milling for cryo-electron tomography

**DOI:** 10.1101/797514

**Authors:** Tobias Zachs, João M. Medeiros, Andreas Schertel, Gregor L. Weiss, Jannik Hugener, Martin Pilhofer

## Abstract

Cryo-electron tomography (cryoET) has become a powerful technique at the interface of structural biology and cell biology, with the unique ability to determine structures of macromolecular complexes in their cellular context. A major limitation of cryoET is its restriction to relatively thin samples. Sample thinning by cryo-focused ion beam (cryoFIB) milling has significantly expanded the range of samples that can be analyzed by cryoET. Unfortunately, cryoFIB milling is low-throughput, time-consuming and manual. Here we report a method for fully automated sequential cryoFIB preparation of high-quality lamellae, including rough milling and polishing. We reproducibly applied this method to eukaryotic and bacterial model organisms, and show that the resulting lamellae are suitable for cryoET imaging and subtomogram averaging. Since our method reduces the time required for lamella preparation and minimizes the need for user input, we envision the technique will render previously inaccessible projects feasible.

## Introduction

Cryo-electron tomography (cryoET) is a powerful imaging technique at the interface of cell biology and structural biology, able to image cells in a near-native state and determine the structure of macromolecular machines in their cellular context (Beck and Baumeister, 2016; Koning et al., 2018; Kooger et al., 2018; Oikonomou and Jensen, 2017; Plitzko et al., 2017). CryoET is restricted to samples that are well below 800 nm in thickness and therefore requires sample thinning techniques for specimens like mammalian cells, *C. elegans*, yeast, cyanobacteria, or biofilms. Biological cryoFIB milling is an emerging sample thinning technique, which uses a Gallium ion beam to ablate segments of the sample in order to generate thin lamellae that can be imaged by cryoET (Marko et al., 2007; Rigort et al., 2010). Unlike previous methodologies, cryoFIB milling produces artifact-free specimens, in which *in situ* structural information is preserved. Its application has led to important insights into mechanisms of cellular function (Albert et al., 2017; Böck et al., 2017; Bykov et al., 2017; Cai et al., 2018; Chaikeeratisak et al., 2019; Delarue et al., 2018; Khanna et al., 2019; Mahamid et al., 2016; Rast et al., 2019; Swulius et al., 2018; Weiss et al., 2019). Unfortunately, however, cryoFIB milling for cryoET is at an early stage of technical maturation and the available techniques are highly manual procedures with relatively low throughput.

In current lamella preparation workflows (Marko et al., 2007; Medeiros et al., 2018; Rigort et al., 2010; Strunk et al., 2012; Zhang et al., 2016), samples are vitrified on transmission electron microscopy (TEM) grids by plunge-freezing. Grids are then transferred to a FIB-scanning electron microscope (SEM) instrument, where potential targets are then identified by SEM and FIB imaging (Supplementary Fig. 1a/b). Using a series of ‘rough milling’ steps, sections above and below the desired lamella are sequentially removed by decreasing the separation between two milling areas and using decreasing FIB milling currents (700 to 100 pA) (Supplementary Fig. 1c-e). Once the lamella is thinned to ∼500 nm, additional targets are identified and thinned by rough milling in a similar manner. To generate lamellae with a final thickness of 100-250 nm, the user returns to each target location and further thins (‘polishes’) each lamella using a low (≤50 pA) current (Supplementary Fig. 1f).

This methodology allows the production of up to 16 lamellae in 10 h (Medeiros et al., 2018), however, during such a session, the process requires constant attention from the operator. The milling process has to be monitored and manual user input is required every 10-15 min, e.g. to execute a series of repetitive tasks such as target identification, positioning milling patterns, changing FIB currents, and visually determining milling end points. This results in a strenuous procedure with a low throughput relative to the time invested by the user, as well as significant idle times due to delays in input from the operator. To overcome these issues, automated sequential cryoFIB milling has become of paramount interest for the field.

### Setup of an automated milling session

Here we report, to our knowledge, the first automated sequential FIB milling method for the preparation of lamellae for subsequent cryoET imaging. Automation was implemented on the Zeiss Crossbeam 550 FIB-SEM instrument, using routines that are available in the SmartFIB software package (Zeiss Microscopy GmbH, Oberkochen, Germany). Particularly important are the modules for stage backlash and drift correction, which are critical for reliable targeting of lamella preparation sites. This allows the user to set up all milling targets and then execute milling in an unattended, fully automated manner.

To begin an automated milling session, FIB current alignments are verified to ensure accurate milling (Fig. 1a). Grids are then loaded into the FIB-SEM instrument. To simplify navigation and target identification, an SEM grid overview image is captured and linked to the stage coordinates as described in the methods. Using the overview image for stage navigation, the first milling site is identified and centered in both the SEM and FIB views (Fig. 1b). To improve the accuracy of mechanical stage movements, the stage is backlash-corrected when moved during automation. To ensure accurate targeting of the milling site, a series of operations is executed before saving the final target position (Fig. 1c). First, stage backlash correction is manually executed and the target is re-centered in the FIB image. Second, the target’s stage coordinates are saved to the stage navigation menu. Third, the stage is manually moved off-target and automatically returned to the saved target location (Fig. 1d). In case the target is *not* properly centered, the above three steps are repeated (Fig. 1e), otherwise the user can proceed.

**Figure 1.**
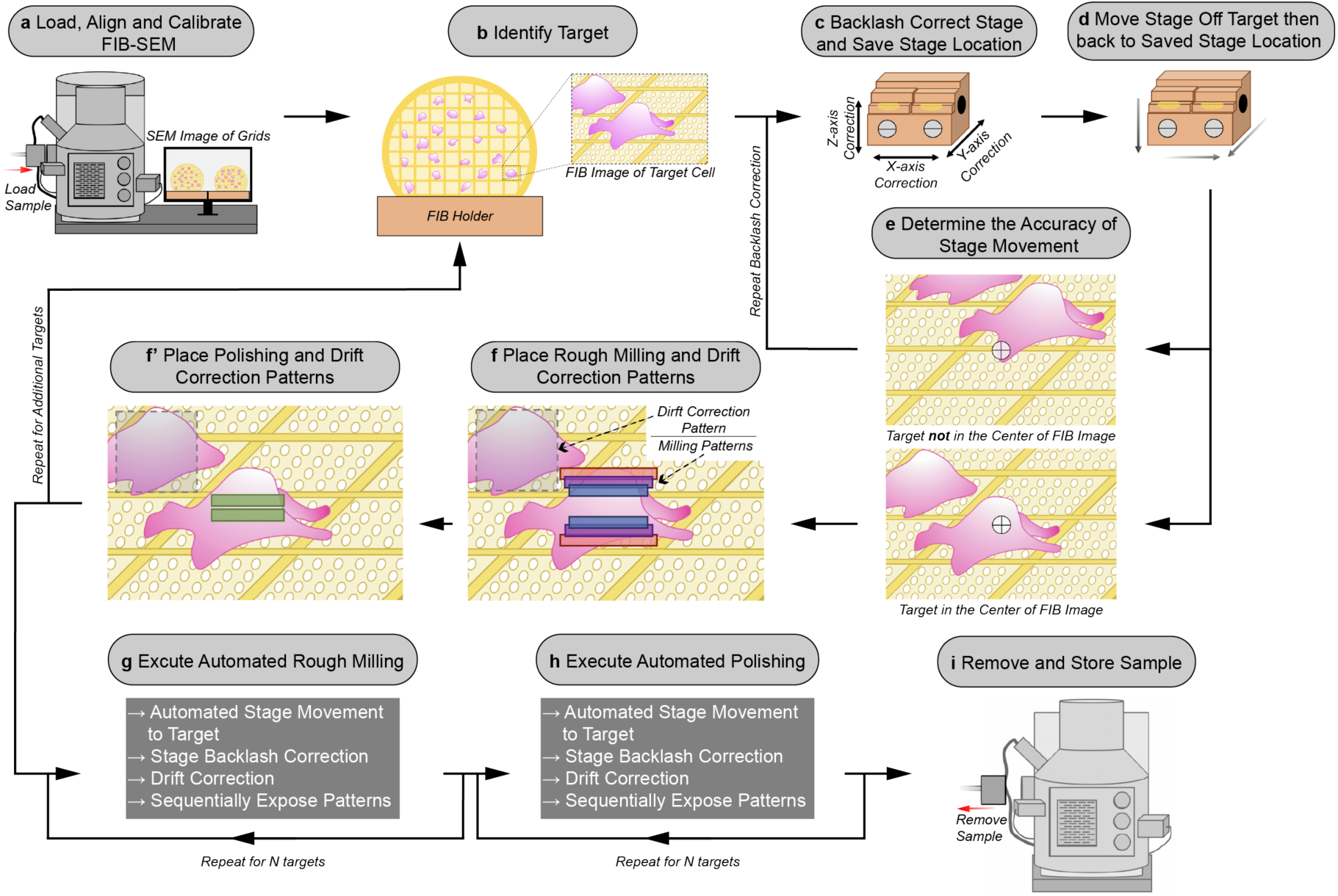
Schematic of the automated sequential cryoFIB milling workflow. **a**: FIB currents are aligned and calibrated, and the sample is loaded into the FIB-SEM instrument. **b**: A target cell is identified on the grid with the FIB. **c**: To correct for errors in mechanical stage movements, backlash correction of the stage is performed. The resulting stage location is saved in the stage navigator. **d**: The stage is randomly moved out of position by the user. Using the saved coordinates, the stage is moved back to the target using the saved position in the stage navigator. **e**: The accuracy of this autonomous stage movement is determined by the user. If the target is not centered in the FIB image, backlash correction is repeated until accurate targeting is achieved (c-e). **f**/**f’**: Rough milling, polishing and drift correction patterns are placed onto the image. Rough milling and polishing patterns are saved separately to the queue. The procedure (b-f’) is repeated to select additional targets. **g**/**h**: Rough milling and lamellae polishing are executed automatically. **i**: The grids with milled lamellae are removed and stored.

Next, patterns with specific currents for rough milling (e.g. 700, 300 and 100 pA) and polishing (e.g. 50 pA) are manually placed onto the FIB image of the target (Fig. 1f/f’). This is achieved by either generating a new set of patterns or by loading previously designed patterns, which is faster and results in more uniform lamellae. To further improve the accuracy of targeting, we incorporated an additional targeting step based on drift correction (Fig. 1f/f’). To implement, each set of milling patterns receives a drift correction box, with user-defined dimensions, which is manually placed in a location close to the target. By capturing and saving an image of the drift correction box, the milling patterns are anchored to their positions on the target.

After saving the first target to the queue, further targets are added by repeating the described procedure. This setup-procedure takes ∼9 min per target.

### Processes during automated milling session

To begin sequential automation, exposure of the rough milling patterns that are saved in the queue is started (Fig. 1g). For each target, the stage automatically moves to the target position and executes stage backlash correction. Next, image shifts are determined between the drift correction image that was recorded during the setup procedure and a drift correction image that is recorded after arriving at the target location. Any existing shifts are compensated for, using FIB beam shifts, to improve the precision of milling. The rough milling patterns are then exposed, from the highest to the lowest current strength. Previously, manual milling methods used a real-time view in order to determine the time that the FIB needs to cut through the specimen. In our automated approach, the exposure time is calculated by the software using a user-specified milling depth (typically 10 µm), milling current, pattern size and material type (e.g. vitrified ice). After exposing the rough milling patterns for the first target, the procedure is automatically repeated for the remaining targets.

Subsequently, the user can decide whether to perform polishing for all targets in a manual or automated manner (Fig. 1h). The automation of polishing follows the routine described above.

### Application of sequential automated milling

During the development of this method, we tested automated sequential milling using the model organisms *S. cerevisiae* strain SK1 (hereafter yeast) and the multicellular cyanobacteria *Anabaena* sp. PCC 7120 (hereafter *Anabaena*) in six independent milling sessions (Table 1). The number of attempted lamellae per session ranged from five to 20 (Fig. 2). Rough milling success, as defined by the presence of a lamella at the targeted location after rough milling, was 99% (n=73). The only failure in lamella production was the result of a user error, as rough milling was accidentally executed on the same target twice (session F). In session B.1 and B.2, lamellae were successfully generated on ten targets that were spread across two grids containing two different samples (Table 1). This shows the robustness of the targeting routine despite sample variations and the execution of large stage movements during automation.

**Table 1.**
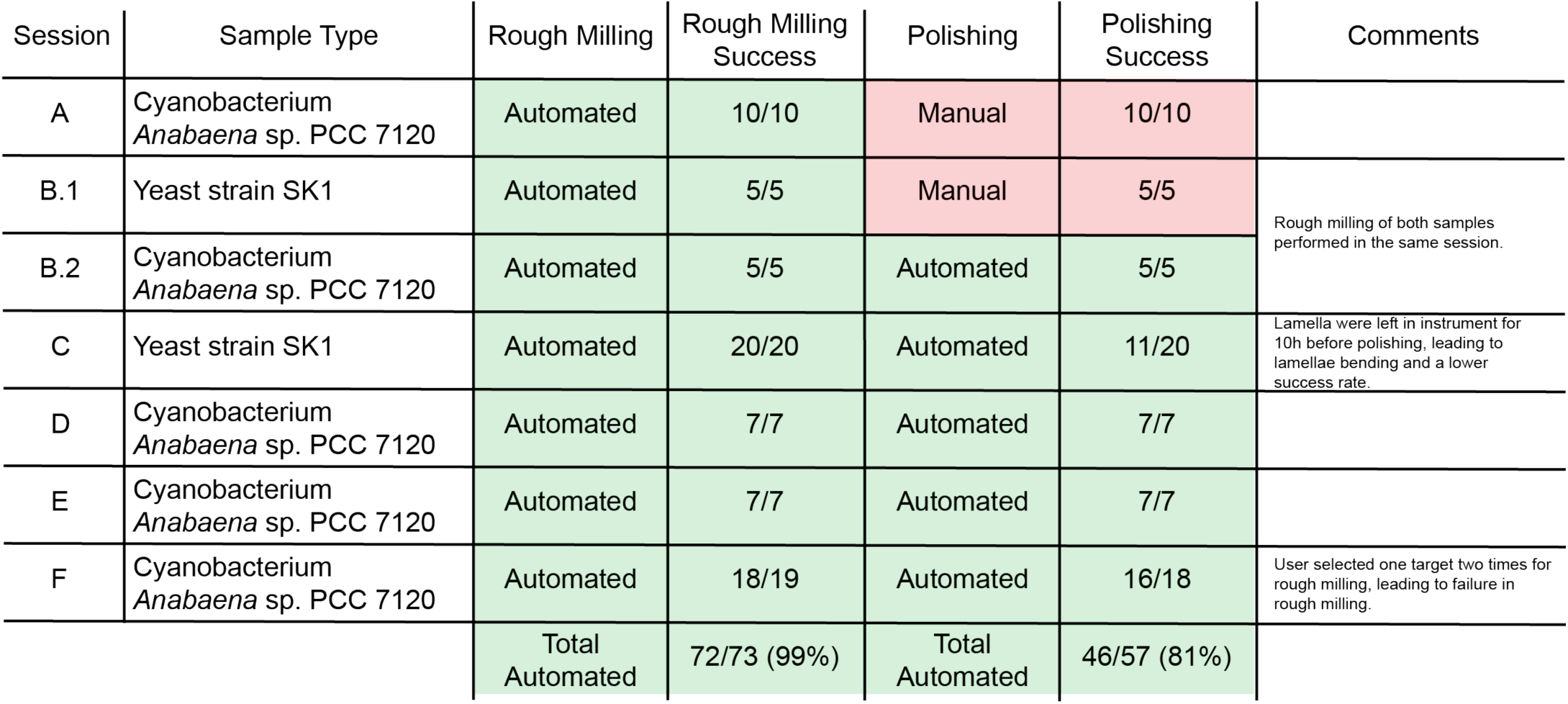
Overview and success rates of milling sessions.

**Figure 2.**
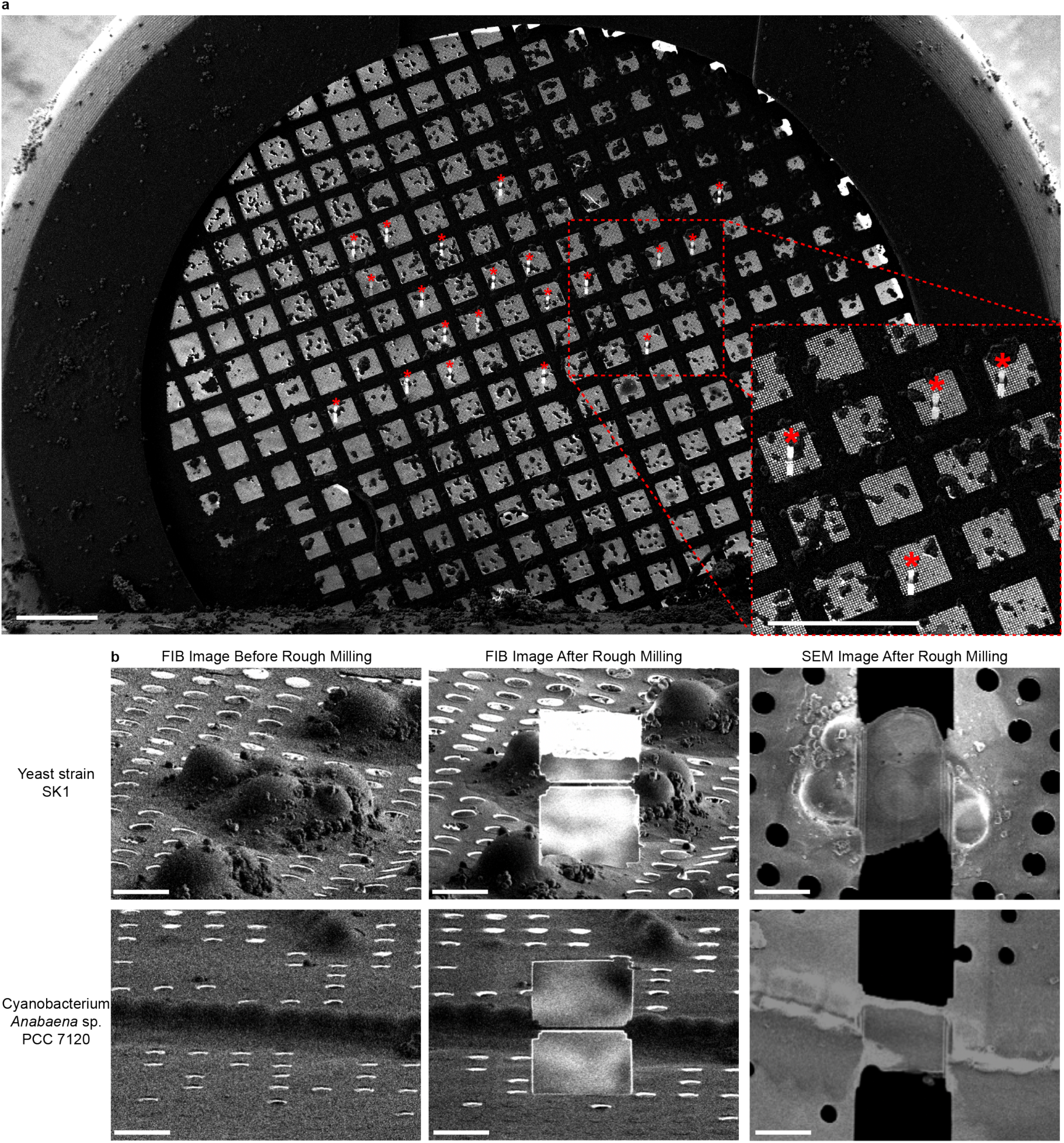
Representative images of lamellae generated by automated sequential rough milling. **a:** SEM grid overview image including 20 yeast targets (asterisks) on which rough milling was performed in an automated sequential manner (session C). Bars, 200 µm. **b:** Representative SEM and FIB images of yeast and cyanobacterial *Anabaena* cells captured before and after fully automated sequential rough milling (session B.1 and B.2). Bars, 5 µm.

While these results present a significant step forward, we next set out to implement automated sequential lamella polishing. In a series of sessions (B.2-F), we milled between five and 20 targets. In total, the success rate (intact lamella detected after polishing) of automated sequential polishing was 81% (n=57 rough lamellae). Importantly, 9 of the 11 failed polishing attempts occurred in session C, in which the rough-milled lamellae were left in the FIB-SEM instrument for 10 h before automated polishing was started. Prior to polishing, these rough lamellae showed signs of bending, which likely resulted in failure in lamellae polishing. Sequential automated lamella polishing should therefore be executed without delay after rough milling. Other reasons for failure in lamella milling could include sample heterogeneity and errors in targeting. If, however, session C were not taken into account, this automated sequential FIB milling methodology would have a 95% (n=37) success rate.

### Assessment of sample quality

In order to assess sample quality, we transferred the grids from all sessions to the cryoTEM. Of the lamellae that were generated in a fully automated manner, 11% (n=46) were lost in transfer. All remaining lamellae could be imaged by cryoET. From the cryotomograms, we determined the lamellae thicknesses to range from 155 to 379 nm (average 232 nm; final polishing patterns were spaced 300 nm apart) (Supplementary Fig. 2). Lamellae that were manually polished (sessions A/B.1) had a comparable average thickness of 258 nm (final polishing patterns were spaced 300 nm apart).

CryoET imaging of the automatically generated lamellae revealed distinct cellular features and macromolecular complexes. Yeast tomograms showed a characteristic nucleoplasm, cytoplasmic ribosomes, nuclear envelope, nuclear pore complexes, and cellular compartments (Fig. 3b). *Anabaena* tomograms showed thylakoid membranes, phycobilisomes and septal junctions (Fig. 3c). To further assess sample and data quality, we performed subtomogram averaging of *Anabaena* septal junctions, which had been characterized recently by a manual cryoFIB milling/cryoET approach (Weiss et al., 2019). From nine lamellae, a total of 412 subvolumes were extracted, averaged and classified in order to remove misaligned particles. The 343 remaining subvolumes were then averaged and symmetrized. The resulting structure revealed key features, including a cap module with five arches, a plug module and a tube module (Fig. 3e-h). Fourier shell correlation (FSC) analyses indicate that the average has a resolution that is similar to a structure that was calculated using the same number of particles extracted from tomograms generated in a previous study (Weiss et al., 2019) (manual milling) (Supplementary Fig. 3).

**Figure 3.**
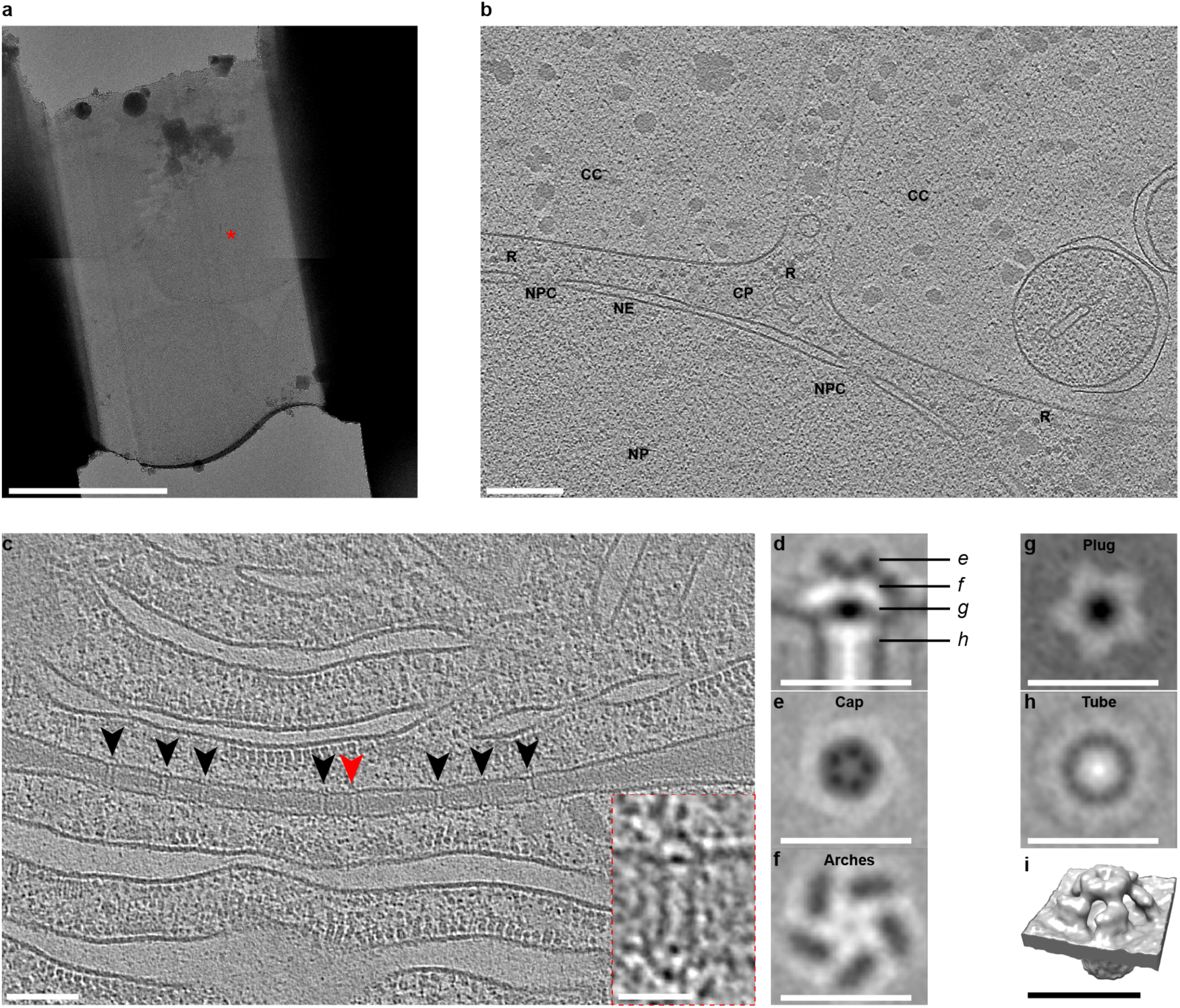
Automated sequential cryoFIB milling results in high-quality lamellae and cryotomograms. **a**: CryoTEM overview image of a typical lamella (session C) containing multiple yeast cells. Red mark indicates the cell imaged in (b). Bar, 5 µm. **b**: Shown is an 18 nm thick slice through a cryotomogram of a yeast cell (session C) [indicated by red mark in (a)]. The thickness of the lamella was determined to be 225 nm. The tomogram shows a characteristic nucleoplasm (NP), nuclear pore complexes (NPC), nuclear envelope (NE), cytoplasm (CP), cytoplasmic ribosomes (R), and other cellular compartments (CC). Bar, 200 nm. **c**: Shown is a 14 nm thick slice through a cryotomogram of a septum between two *Anabaena* sp. PCC 7120 cyanobacteria cells (session F). The thickness of the lamella was determined to be 208 nm. Arrowheads indicate septal junctions. The inset shows a magnified view of the septal junction indicated by a red arrowhead. Other cellular features are cytoplasmic membranes (CM), phycobilisomes (PB), thylakoid membranes (TM) and septal peptidoglycan (PG). Bars, 100 nm and 25 nm (inset). **d-i**: A subtomogram average was generated by 5-fold symmetrizing 343 septal junctions that were extracted from nine tomograms. Shown are longitudinal (d) and perpendicular (e-h) slices (thickness 0.68 nm) and a surface rendering (i) of the symmetrized average. The observed characteristic structural modules were similar to a recent study that applied manual cryoFIB milling (Weiss et al., 2019) (also see Supplementary Fig. 3). Bars, 25 nm.

## Discussion

In conclusion, our automated sequential cryoFIB milling method allows for the production of high-quality lamellae for cryoET imaging and will impact cryoFIB/cryoET projects in several ways. First, the time investment by the operator is significantly reduced from ∼10 h in a manual milling session to ∼2.4 h for an automated sequential milling session, assuming 16 targets are milled. Second, by removing the need for frequent user inputs and idle times, the minimum required machine time is reduced from ∼38 min (Medeiros et al., 2018) (i.e. 16 lamellae in 10 h) to ∼25.5 min (9 min setup plus 16.5 min milling) per lamella. Third, based on the robustness and customizable nature of the method, the procedure can be adapted to a wide range of samples and milling techniques (Toro-Nahuelpan et al.; Wolff et al., 2019). Fourth, the automated procedure will allow the user to systematically explore novel milling methods by reusing uniform milling patterns. Fifth, the method can generally be combined with correlated approaches that allow for target pre-screening, for instance cryo-light microscopy or cryo-FIB-SEM volume imaging (Eibauer et al., 2012; Gorelick et al., 2019; Koning et al., 2014; Schertel et al., 2013; Schorb et al., 2017; Sviben et al., 2016; Vidavsky et al., 2016). That said, the higher throughput achieved by automated cryoFIB milling (shown here) in combination with fast cryoET data collection schemes (Chreifi et al., 2019; Eisenstein et al., 2019), might in many cases eliminate the need for target pre-identification by correlated approaches. Altogether, the development of automated sequential cryoFIB milling renders cryoET applicable to previously unfeasible projects.

## Acknowledgments

Joao Matos is acknowledged for providing resources for culturing yeast cells. We thank Saskia Mimietz-Oeckler and Andreas Hallady (Leica Microsystems GmbH, Vienna, Austria) for technical support. We thank ScopeM for instrument access at ETH Zürich. MP was supported by the Swiss National Science Foundation (#31003A_179255), the European Research Council (#679209) and the Nomis Foundation.

## Methods

### Overview of the equipment and workflow

The method was established and tested on a Crossbeam 550 FIB-SEM instrument (Carl Zeiss Microscopy) equipped with a copper band-cooled mechanical cryo-stage and an integrated VCT500 vacuum transfer system (Leica Microsystems). The detectors used included an InLens secondary elelctron (SE) detector for determining grid topology (Carl Zeiss Microscopy) and a SE2 detector for identifying milling targets and assessing the ice thickness (Carl Zeiss Microscopy). In our workflow, EM grids were prepared with budding yeast strain SK1 and *Anabaena* sp. PCC 7120, and clipped into FIB milling Autogrids (ThermoFisher Scientific, Waltham, Massachusetts, U.S.). These grids were then mounted onto a pre-tilted Autogrid holder (Medeiros et al., 2018) (Leica Microsystems) using a VCM loading station (Leica Microsystems). Using the VCT500 shuttle, the Autogrid holder was transferred to an ACE600 cryo-sputter coater (Leica Microsystems) under cryogenic conditions and the samples were sputter-coated with a 4 nm thick layer of tungsten. After sputter coating, the samples were transferred into the Crossbeam 550 using the VCT500 shuttle. In the Crossbeam 550, the gas injection system (UniGIS) was used to deposit an organometallic platinum precursor layer onto each grid. Automated sequential FIB milling was subsequently set up and executed. Sample preparation, plunge-freezing, Autogrid mounting, holder loading and vacuum cryo-transfer steps were executed similarly to what was described in Medeiros et al. 2018. Any deviations to the previously published protocol are described below.

### Cell culture and plunge freezing

FIB milling tests were performed using the cyanobacterial strain *Anabaena* sp. PCC 7120 and the *S. cerevisiae* strain SK1. The *Anabaena* strain was grown and prepared for FIB milling as previously described in Weiss et al. 2019. Yeast cells were prepared as previously described by Medeiros et al. 2018.

### Equipment calibration

To ensure that automated sequential FIB milling was successful, the Crossbeam 550 was properly aligned. While the SEM column alignments are stable and non-essential during automated milling, the FIB alignment between different currents at a given voltage (30 kV for biological cryo-samples) should be checked and optimized. Typically, this calibration is done weekly or when deemed necessary and takes roughly 60 min to complete. In case of deviation, on-the-fly adjustments are possible on a loaded cryo-sample, however, standard calibration procedures are best performed on a silicon wafer due to its structural homogeneity, which allows better evaluation of the FIB beam shape. Once inserted into the chamber, the stage was tilted by 54 ° to be perpendicular to the FIB beam and then moved to the working distance (i.e. coincidence point). Using the ‘spot’ function in an unexposed sample region, the beam was focused to its spot size allowing it to burn a hole into the silicon. If the current is properly calibrated, then the beam will produce a spot that is round with sharp edges. This was best seen when using a mixed signal of the InLens and SE2 detector. If a beam spot had imperfections, like a tailing edge, beam parameters including focus, stigmatism and aperture alignments need to be improved and saved. After optimizing these parameters for each current, all currents were aligned against the reference current. This was best performed by centering an easily recognizable structure like a burnt hole for each beam onto the exact position in the image taken with the reference current. Finally, to ensure that the currents were properly aligned, a location is imaged by each current. If properly aligned, switching between currents should not lead to focus changes or beam offsets.

### Sample coating

To enhance sample conductivity and decrease the effects of charging, EM grids were coated with a ∼4 nm layer of tungsten using the sputter coating head on the ACE600. After inserting the holder into the FIB-SEM, a protection layer of organometallic platinum precursor was deposited onto each grid to minimize the curtaining effect. For cold deposition of platinum precursor, the holder was moved 3 mm below the coincidence point and was tilted to 20 degrees. By positioning the gas injection system (GIS) needle above each grid and opening the GIS for 30 s, a layer of platinum precursor was deposited onto the sample. Since the GIS needle was mounted at a similar angle as the FIB column, deposition of platinum occurred preferentially on the side of the cells where the FIB beam hits the sample, ensuring the best protection. For deposition under cryo-conditions, it is essential that the heating element of the GIS needle and reservoir are turned off to keep the system at room temperature (28 °C).

### Stage registration

To assist in the identification of targets, overview images of an entire EM grid are taken. On the Zeiss Crossbeam 550, these images can be coupled to the stage navigation. To calibrate stage registration a high-resolution (4096 x 3072 pixels, 35x) overview image was taken with the SE2 detector, which provided the best information for identifying targets inside the vitrified ice and determining ice thickness. This overview image was then loaded onto the stage navigation bar and registered by correlating three distinctive points on the image to their specific positions on the stage as observed in the live SEM view. After completion, double clicking on a desired target image location in the navigation bar automatically moves the stage to the location of interest. In addition, backlash correction was also included for all automated stage movements, using the user preference settings of the software SmartSEM (Carl Zeiss Microscopy).

### Defining milling materials

To permit unsupervised automation of lamellae production, the Crossbeam 550 was calibrated to mill a cross-section with a specified depth through the sample. To ensure proper milling, the system needs to be calibrated for a distinct ‘material’ so that the correct milling parameters like dose are applied during milling. For cryo-TEM lamella preparation the material “vitrified ice” was created using a dose calibration of 20 mC / cm^2^ being equivalent to a milling depth of 1 µm in cross-section mode.

### Parameters for imaging and milling

For SEM imaging, voltages from 1.9 - 5 kV and a constant current of 28 pA were used. To capture SEM images, we most commonly used the InLens detector to obtain surface information of the sample. During FIB imaging, on the other hand, a fixed voltage of 30 kV and a low current (20pA) was used. FIB images were usually captured by using the SE2 detector, which is less sensitive to imaging-induced charging. During automated sequential milling, four sets of currents above and below the desired lamellae were used. For rough milling 700 pA, 300 pA and 100 pA currents were implemented. To then polish the lamellae, a 50 pA current was used. For milling, we defined the patterns to be executed using bi-directional and cross-section cycle mode with a 10 µm milling depth.

### Automated sequential FIB milling protocol

To generate high quality lamellae, it was essential to prepare the FIB-SEM and sample for automated sequential milling. Preparations included checking and calibrating the FIB currents, coating the sample with a layer of tungsten and organometallic platinum, and performing stage registration. Once these steps were executed, automated sequential milling was initiated by identifying and setting up milling targets.

The grid overview image in the stage navigator was used to identify a milling target. The identified target was then manually centered in the live FIB view with the aid of the SEM. To improve the accuracy of automated stage movements, backlash correction was performed manually and implemented for all automated stage movements. The target’s stage coordinates were then saved in the stage navigator. To ensure that the instrument was able to perform targeting during automation, the stage was manually moved away from the target and then instructed to move back to its saved location. The target was located using the live FIB view and if necessary, manually centered again. If manual centering was required, the new stage location was saved and the instrument’s ability to perform targeting was tested again. To ensure successful milling during automation, it was essential to refine the stage location until the stage was able to perform targeting successfully.

Once an accurate stage movement was achieved, milling patterns were placed onto a target FIB image captured using SmartFIB. In SmartFIB, each pattern contains specific milling conditions (i.e. current, milling depth, size, shape, etc.) and a designated FIB milling location. SmartFIB allows the placing of multiple patterns with different conditions onto a single FIB image in order to perform automated milling. Patterns were placed and their properties were changed by using the SmartFIB GUI in the ‘Attributes’ tab. When testing this methodology, we placed eight rectangular milling patterns: six rough milling and two polishing patterns (Supplementary table 1). The final polishing patterns were spaced 300 nm apart, which from our experience results in an average lamella thickness of 225-275 nm. To make uniform lamellae it was also possible to save these eight patterns as a recipe, which can be dragged and dropped onto images of other milling targets. To then save these milling patterns, it was essential to separate the rough and polishing patterns. This was accomplished by deleting the polishing patterns from our recipe, saving only the rough milling patterns, undoing the deletion of the polishing patterns (using the SmartFIB Undo button), deleting all rough milling patterns and then saving only the polishing patterns.

To improve the targeting accuracy of this methodology, a drift correction step was also added to each set of rough and polishing milling patterns immediately before being saved. This was done in the SmartFIB ‘attributes’ tab, by capturing and saving an image of a defined region of the FIB view. During the automated protocol SmartFIB would use this image to perform image recognition before beginning milling and compensate for small shifts to ensure the milling patterns are placed correctly on the target. When testing this methodology, it was important to save and then load the same drift correction image for both the rough and polishing milling patterns. This ensured the highest accuracy when moving from rough milling to polishing.

After saving a set of rough and polishing patterns, the described method can be repeated for further targets. For an automated protocol, about 9 min were needed to set up each target. It is possible to also automate the milling of targets found on separate EM grids. Once satisfied with the number of targets, all rough milling recipes in the SmartFIB queue were selected and exposed. Exposure of a typical rough milling target takes about 12 min. Upon completion, rough milling targets were observed using the SEM and FIB to determine their quality. To then initiate polishing, it is possible to either tick all polishing recipes and expose them, or individually move to each target using SmartFIB, take a FIB image, manually drag polishing patterns into place and expose the lamella. Polishing typically took about 4.5 min. Once all targets are polished, the lamellae are removed from the instrument and stored. It is essential to note that we aimed to keep the lamellae in the instrument for <2 h after beginning polishing to minimize contamination. In theory, this limits our lamellae production to ≤20 targets. If, however, aspects including the milling depth, pattern sizes or currents were changed, it would be possible to generate more lamellae. Note that in our attempts, grids with milled lamellae were transported in a dry-shipper from Zeiss Oberkochen, Germany to Zürich, Switzerland prior to cryoET imaging, possibly resulting in some lamellae breaking. An overview of all the milling attempts performed can be found in Table 1.

### Cryo-electron tomography, tomogram reconstruction and subtomogram averaging

Data was collected on a Titan Krios 300kV electron microscope (ThermoFisher) equipped with a field emission gun, imaging filter (Gatan, Munich, Germany) (slit width 20 eV) and K2 Summit direct electron detector (Gatan). To generate an overview of each grid, grid montages were collected at 135x magnification using SerialEM (Mastronarde, 2005). UCSF Tomo (Zheng et al., 2007) was used for automated recording of tilt series (+60° and −60° tilt range, 2° increments). Data was collected at a defocus of −8 µm, total accumulated dose of ∼140 e^-^ / Å^2^ and pixel size of 3.38 Å. Tomogram reconstruction and subtomogram averaging was performed according to Weiss et al. 2019. Briefly, tomograms were reconstructed using the IMOD package (Kremer et al., 1996) and subtomogram averaging was performed using PEET (Nicastro et al., 2006). A total of 412 particles were extracted and averaged in a box of 44 × 44 × 44 pixels with a pixel size of 0.68 nm. PEET classification was then used to remove misaligned particles (343 final particles). 5-fold symmetry was applied to obtain the final average. The FSC (Fourier Shell Correlation) was generated by using the PEET command calcFSC.

## Data and code availability

Example tomograms of yeast and *Anabaena* lamellae milled in a fully automated manner and the final septal junction subtomogram average determined in this study were deposited to the Electron Microscopy Data Bank (accession number EMDB xxx-yyy for the tomograms and zzz for the subtomogram average).

---

**Supplementary Figure 1.**
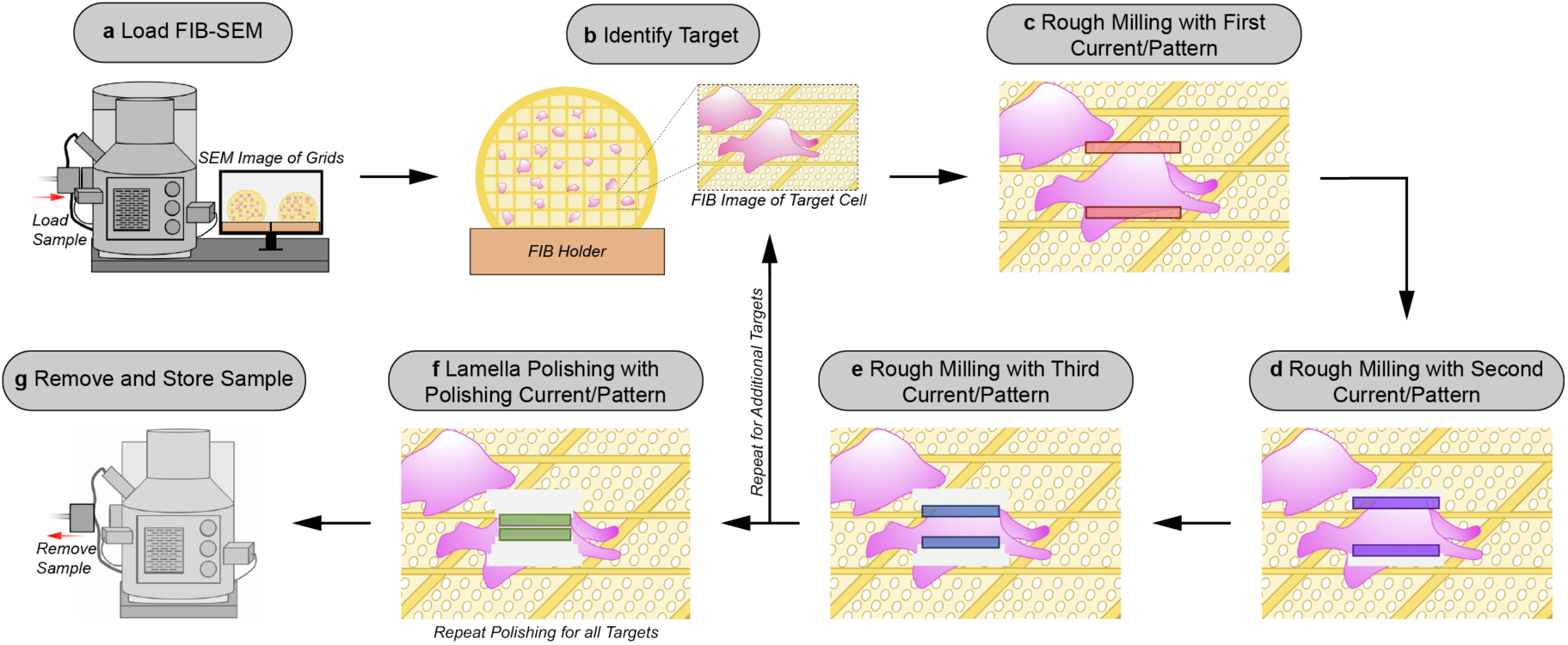
Schematic of the manual cryoFIB milling workflow. **a:** The sample is loaded into the FIB-SEM instrument. **b:** A target is centered in the FIB image. **c-e:** The first pair of rough milling patterns is placed on the target and milling is executed (c). Lamellae milling is observed via a live FIB view to determine when a milling step is completed. The same procedure is repeated for the second (d) and third (e) rough milling patterns. After rough milling of the target is completed, additional targets can be milled by repeating steps b-e. **f:** Rough-milled lamellae are polished with a fourth set of milling patterns. Polishing is repeated for all rough-milled lamellae. **g:** The grids with milled lamellae are removed and stored.

**Supplementary Figure 2.**
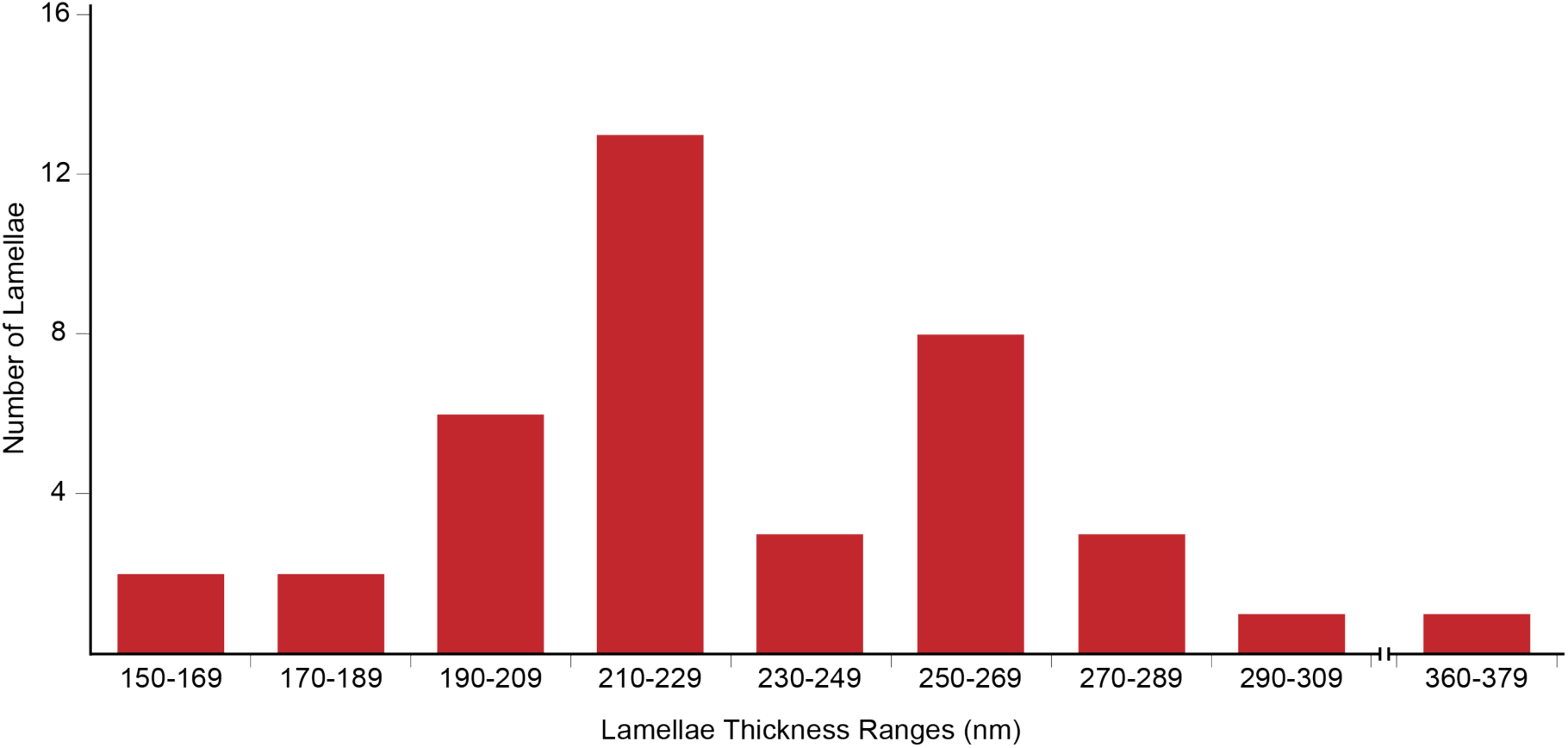
Distribution of lamellae thickness. The plot shows the distribution of the thickness values of fully automated sequential FIB-milled lamellae, as determined by cryoET imaging. The final polishing milling patterns were spaced 300 nm apart.

**Supplementary Figure 3.**
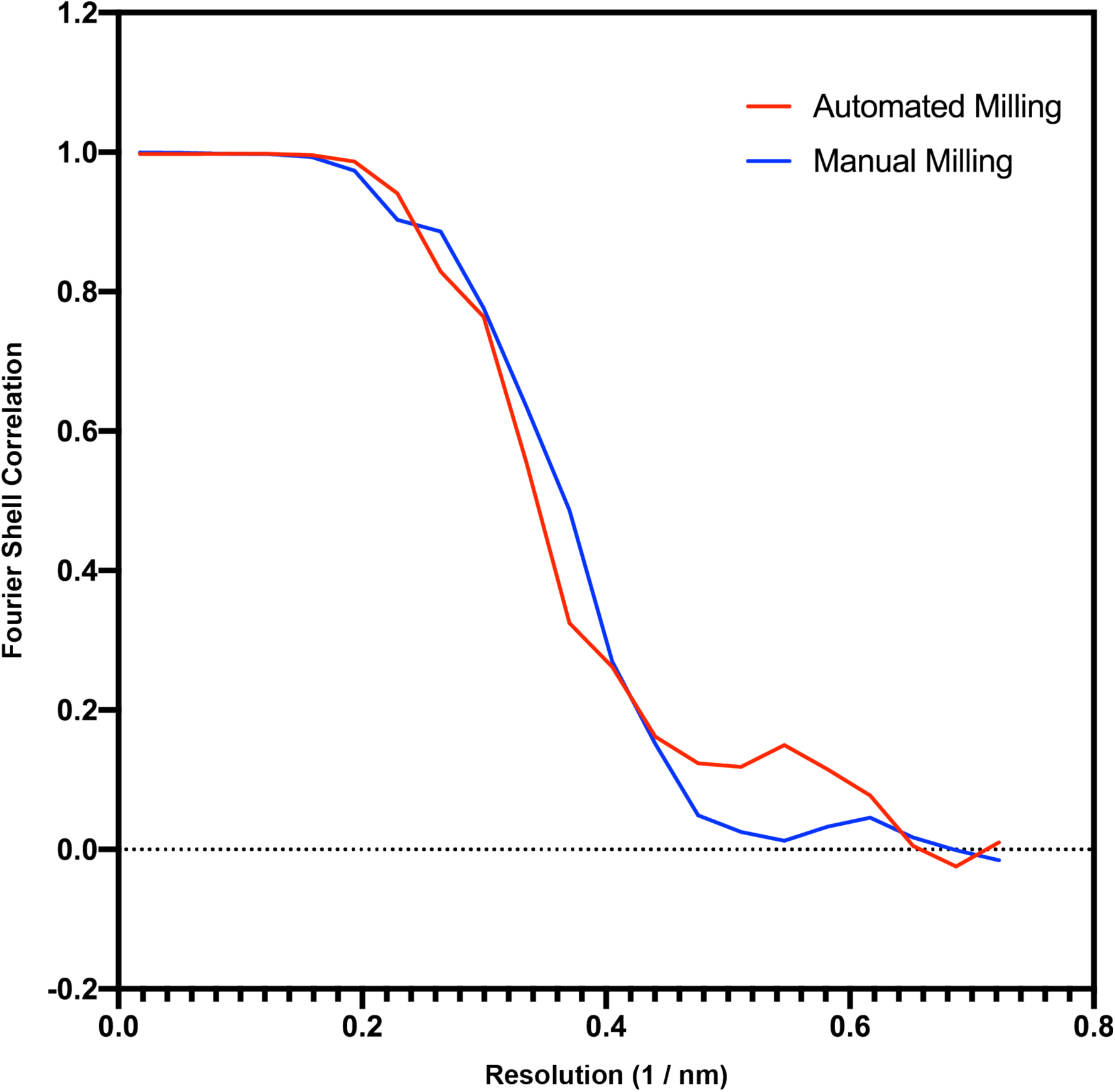
Comparison of data quality between manual and automated milling. Shown is a Fourier Shell Correlation (FSC) curve (blue) for the septal junction subtomogram average shown in Figure 3i (resulting from automated milling). The second curve (red) results from a dataset published previously (Weiss et al., 2019) (resulting from manual milling) and was calculated with the same number of randomly selected subvolumes after 5-fold symmetrization (n=1715). Both approaches result in a comparable resolution estimate.

**Supplementary Table 1.** Dimensions and currents used for each milling pattern during automated sequential lamellae preparation.

|  |  |
| --- | --- |
| Milling Patterns 1 and 2<br>(Rough Milling 1) | 30 kV 700 pA<br>9 x 5 $\mu\text{m}$ rectange<br>Patterns spaced 2 $\mu\text{m}$ apart |
| Milling Patterns 3 and 4<br>(Rough Milling 2) | 30 kV 300 pA<br>8 x 2 $\mu\text{m}$ rectange<br>Patterns spaced 1 $\mu\text{m}$ apart |
| Milling Patterns 5 and 6<br>(Rough Milling 3) | 30 kV 100 pA<br>7.5 x 1 $\mu\text{m}$ rectange<br>Patterns spaced 500 nm apart |
| Milling Patterns 7 and 8<br>(Lamella Polishing) | 30 kV 50 pA<br>7 x 0.5 $\mu\text{m}$ rectange<br>Patterns spaced 300 nm apart |
| Drift Correction Pattern<br>(Rough Milling and Polishing) | 3 x 3 $\mu\text{m}$ rectangle |

